# Fixation Drift Increases as a Function of Time-on-Task

**DOI:** 10.1101/2024.09.05.611445

**Authors:** Lee Friedman, Oleg V. Komogortsev

**Affiliations:** Department of Computer Science, Texas State University, San Marcos, Texas, USA

## Abstract

Ocular fixations contain microsaccades, drift and tremor. We report an increase in the slope of linear fixation drift as a function of time-on-task (TOT). We employed a very large dataset (322 distinct subjects, multiple visits per subject). Subjects performed a random saccade task. The task, in which the target dot jumped randomly over the display area every 1 sec, was 100 sec in duration. Fixations were identified using a published classification method. For each fixation, we regressed eye position against time across multiple segment lengths (50, 100, 200, 300, 400, and 500 ms). We started with the first sample and continued until no further regressions were possible based on the particular segment length being evaluated. For each segment length, each fixation was characterized by a single value: the maximum slope over the segment length. The slopes were expressed in deg/sec. We were not interested in the direction of the linear drift so we took the absolute value of the slope as the measure. For data analysis, each 100 sec task was divided into five 20 sec epochs. We found that median slope increased across epochs in both session recordings. Although similar trends were found regardless of segment length, the results were clearer and more consistent when using segment lengths of 200 ms or greater. Although we describe these changes in linear drift as related to time-on-task (TOT), we think it is likely, though no certain, that these effects are due to some sort of short-term oculomotor fatigue.

## Introduction

It is well established that during “ocular fixation” the eye is not still. There are three types of eye-movements that occur: microsaccades, drift and tremor [1]. According to Martinez-Conde and Macknik [2]:

> Drift is the slow(usually less than 2 *deg s*^−1^), curvy motion that occurs during the periods of relative fixation between microsaccades and/or saccades. Referred to as ‘slow control’ in some early studies, drift is typically not conjugate and it resembles a random walk.

It has been known since the early 1800’s that deliberately focusing one’s gaze on a target can make unmoving images in the surrounding region gradually fade away. This type of vanishing illusion is known as Troxler fading [3]. According to [2]:

> Recent research indicates that whereas only microsaccades effectively restore visibility after perceptual fading during fixation, both microsaccades and drift serve to prevent fading from occurring in the first place.

DiStasi et al. [4] is the first, and as far as we are aware, only publication to report that fixation drift increases as a function of fatigue. In that study, subjects performed a simplified air traffic control task. The entire task took two hours to complete and consisted of four 30 minute blocks. Drift periods were defined as the eye-position epochs between microsaccades, saccades, overshoots and blinks. Drifts shorter than 200 ms were discarded. For each drift, the key measures were mean and peak velocity. These authors noted a statistically significant linearly increasing drift mean velocity and peak velocity from the first 30 min block to the last 30 min block. The authors conclude that increased fixation instability occurs with time on task (TOT).

Although that prior study [4] stimulated our interest in this topic, our study is distinctive in a number of important ways: (1) Di Stasi et al. [4] studied the effect of TOT on drift velocity over a 2 hour period whereas our study evaluated changes in drift related to TOT where the entire task was completed in 100 seconds. (2) Di Stasi et al [4] evaluated drift during the performance of a simplified air traffic control task, we evaluated drift during a random saccade task. (3) We defined drift as linear drift only, i.e., we estimated the slope of fixation drift over several segment lengths (50, 100, 200, 300 and 400 ms). (3) The [4] study was based on the evaluation of twelve subjects whereas our study included data from 322 subjects. Since subjects were evaluated in two sessions over multiple rounds, our results are based on 1,762 separate eye movement recordings sessions. Despite these clear differences between the the Di Stasi [4] study and our study, both found evidence for increased fixation drift as a function of TOT.

## Materials and methods

### The Eye Tracking Database

The eye tracking database employed in this study is fully described in [5] and is labelled “GazeBase” It is publicly available (https://figshare.com/articles/dataset/GazeBase_Data_Repository/12912257). The data were accessed on April 12, 2024. The authors had no access to information that could identify individual participants during or after data collection.

All details regarding the overall design of the study, subject recruitment, tasks and stimuli descriptions, calibration efforts, and eye tracking equipment are presented there. There were 9 temporally distinct “rounds” over a period of 37 months, and round 1 had the largest sample (see Table 1). For round 1, there were 151 females and 171 males. Subjects in each successive round needed to have been present in all preceding rounds. This report is based on all available data, including subjects from rounds 1 to 9. There were 881 subject visits (Table 1) and two sessions (*≈* 20 min apart) per visit for for a total of 1,762 studies or recordings. Briefly, subjects were initially recruited from the undergraduate student population at Texas State University through email and targeted in-class announcements. Each session consisted of seven tasks. The only task employed in the present study was the random saccade task. During the random saccade task, subjects were to follow a white target on a dark screen as the target was displaced at random locations across the display monitor, ranging from ± 15 and ± 9 of degrees of visual angle (dva) in the horizontal and vertical directions, respectively. The minimum amplitude between adjacent target displacements was 2 dva. At each target location, the target was stationary for 1 sec. There were 100 fixations per task (100,000 samples per task). The target positions were randomized for each recording. The distribution of target locations was chosen to ensure uniform coverage across the display. Monocular (left) eye movements were captured at a 1,000 Hz sampling rate using an EyeLink 1000 eye tracker (SR Research, Ottawa, Ontario, Canada).

**Table 1.** Number of Subjects per Round.

| Rounds | Subjects |
| --- | --- |
| 1 | 322 |
| 2 | 136 |
| 3 | 105 |
| 4 | 101 |
| 5 | 78 |
| 6 | 59 |
| 7 | 35 |
| 8 | 31 |
| 9 | 14 |
| Sum | 881 |

The gaze position signals for the random saccade task were classified into fixations, saccades, post-saccadic oscillations (PSOs) and various forms of artifact, using an updated version of our previously reported eye-movement classification method [6]. Although the code for actual saccade detection is complex, we describe it here in a very brief format. Saccade detection was based on an analysis of smoothed radial velocity. To detect saccades, the first step was to look for an event with a radial velocity above 100 deg/sec. The start and end of the saccade were based local velocity minima preceding and following the peak radial velocity of the event.

### Screening Fixations

#### Removing fixations adjacent to artifact

Any fixation which was immediately preceded or followed by any type of artifact (e.g., blink artifact), was excluded from this study.

#### Removal of fixations that are part of Square Wave Jerks

According to [7], one type of saccadic intrusion, “Square Wave Jerks” (SWJ) are:

> “…small (typically 0.5 degree), horizontal, involuntary saccades that take the eyes off the target and are followed, after an intersaccadic interval of about 250 milliseconds, by a corrective saccade that brings the eyes back to the target. They may occur in normal individuals at frequencies of 20 per minute or greater.” Page 250.

Since fixations during these SWJ are fundamentally different from other fixation types, we wanted to exclude them. To this end, we develop a MATLAB (Natick, Massachusetts) script to detect SWJ and remove the fixations associated with SWJ from our dataset. To illustrate the results, for round 1 only, we found a total of 1,467 SWJ. Of 322 subjects, 54 had no SWJs. Of the remaining 268 subjects, 55 had only 1 SWJ. The median number of SWJ per subject was 2.5 (25th percentile = 1, 75th percentile= 5 SWJ per subject). One subject had 26 SWJ in session 1 and 22 SWJ in session 2. Our MATLAB script for detecting SWJ, 61 eye movement datasets, and 89 example images of SWJ are available online at https://hdl.handle.net/10877/18499.

### Calculation of Fixation Drift

As illustrated in Fig. 1 (A), we start with the position data (either horizontal or vertical). This particular fixation is 715 samples long. In the illustrated case, we are fitting a line that is 300 ms in length. (We also tested, and will report on, other lengths (50, 100, 200 msec lengths)). For this particular analysis, the fixation has to be at least 301 ms in duration. Shorter fixations were not analyzed. The first step is to regress the position signal, starting at sample 1 of each fixation, against a vector from 1 to 300 in steps of 1. Because we were interested in quantifying drift regardless of direction, we take the absolute value of the slope. The raw slope is expressed in degrees per ms, so we multiply the slope by 1000 to get degrees per second. Once we have the slope starting at point 1, we then proceed to the next sample and repeat the analysis. This keeps going until the last sample that allows for a test of a 300 ms linear fit. In Fig. 1 (B), we plot the 416 slopes for this fixation. The measurement of drift is taken as the largest such slope for each fixation.

**Fig 1.**
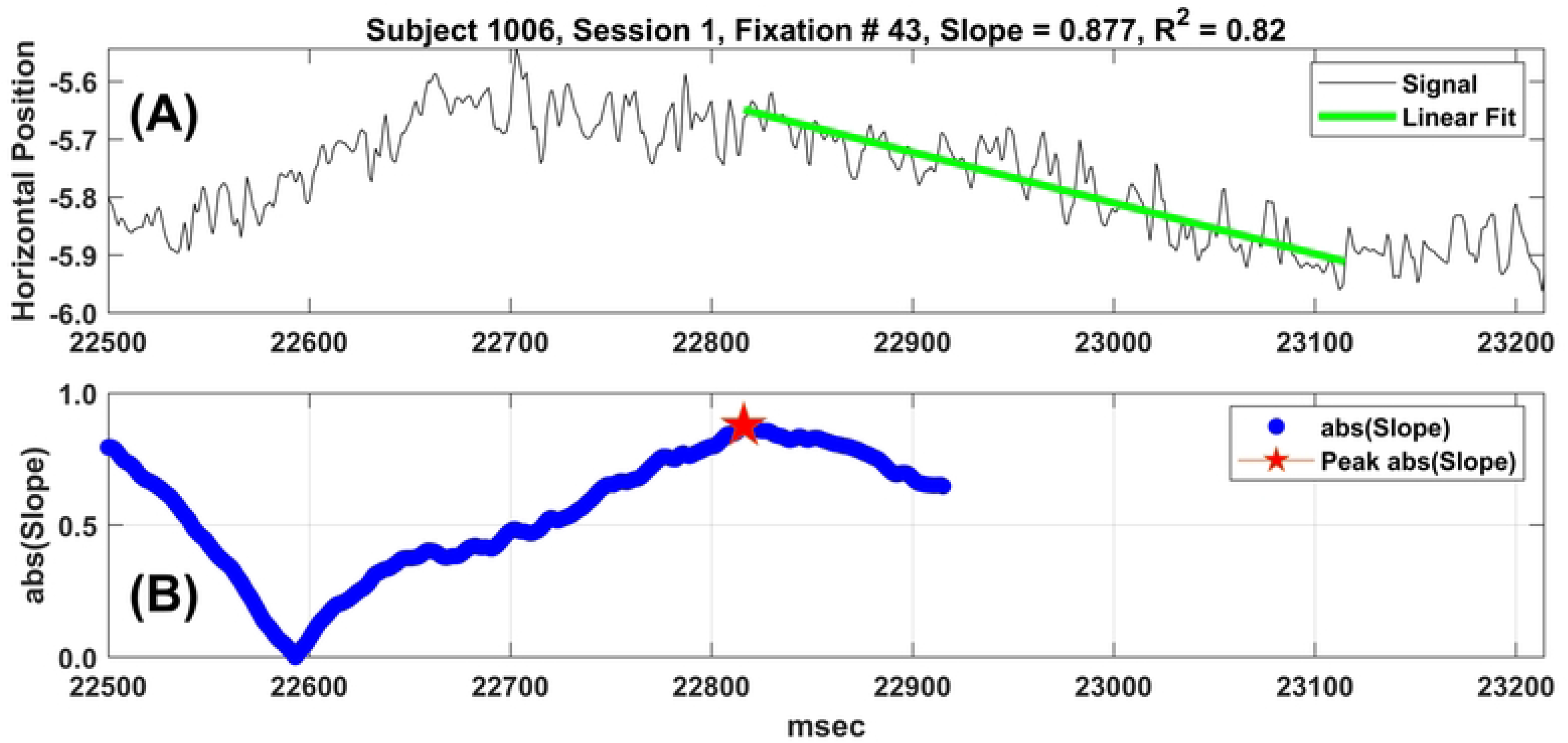
Illustration of the Fixation Drift Evaluation Method. (A) An example fixation from subject 1006, session 1. This is fixation number 43. It is 715 ms in length. The horizontal position is displayed. (B) This is a plot of 416 slopes. Starting at the first point, we regressed the first 300 eye-position data points onto a vector of ms numbers from 1:300. The slope of this regression represents the slope of eye position over this interval. Since we were interested in quantifying drift regardless of direction we take the absolute value of the slope. The raw slope expresses the drift in degrees per ms, so we multiply the slope by 1000 to obtain the slope per second. The red star is the point with the highest slope (absolute value). The green line in (A) is the fitted regression line plotted on the horizontal position signal starting at the point of maximum slope.

### Statistical Analyses

#### Analysis of Linear Trend of Median Slope across Epochs

For each session, each random saccade task was 100,000 msec long. For each session the 100,000 ms were divided up into five 20,000 ms “epochs”. For each epoch we computed the median slope across all available fixations. If a fixation started during one of the five epochs, the fixation was assigned to that epoch. This resulted in 10 medians across 10 epochs, five for session 1 and five for session 2. After viewing the results and noticing an increasing median slope from epoch to epoch, we regressed these 10 medians across all 10 epochs, renumbering epochs for session two to 6 thru 10. For each segment length, this produced a median slope, an *R*^2^ for the linear relationship between slope and epoch, and an F-value and p-value for the regression. The error degrees of freedom (df) for all of these analyses was 8.

#### Statistical Comparison of Slope Estimates across Epochs

A Kruskal-Wallis (KW) test was conducted separately for each segment length and position direction (horizontal or vertical). Each KW test produced a *χ*^2^, a numerator *df*, a denominator *df* and a *p* − *value*. Since all of these *χ*^2^ values were statistically significant, each test was followed by a multiple comparison procedure using Tukey’s Honestly Significant Difference (HSD) method with a *p* − *value*, corrected for multiple comparisons, of 0.05.

## Results

### Comparison of slope results across various segment lengths

We analyzed six different segment lengths (50, 100, 200, 300, 400 and 500 ms). Table 2 presents the median analysis for horizontal position for all segment lengths. As expected, the median slope decreased as the segment length increased. The pattern of results for each segment length were similar. The highest *R*^2^ was for a segment length of 300 msec (0.986). To keep the number of figures manageable, we only present figures below based on a segment length of 300 ms, although figures for all segment lengths are available at (https://hdl.handle.net/10877/19260). The results for vertical signals is presented in Table 3. In this case, the highest *R*^2^ was for a segment length of 400 msec (0.987), although the *R*^2^ for a segment length of 300 msec was very similar (0.985). The median slopes for horizontal and vertical signals were similar across segment lengths.

**Table 2.** Median Fits by Segment Length - Horizontal Signals.

| Segment Length (ms) | Median Slope <sup>2</sup> | $R^2$ | F-value<sup>1</sup> | p-value |
| --- | --- | --- | --- | --- |
| 50 | 3.10 | 0.904 | 75.6 | 0.0001 |
| 100 | 1.75 | 0.957 | 177.4 | 0.0001 |
| 200 | 1.10 | 0.981 | 410.4 | 0.0001 |
| 300 | 0.84 | 0.986 | 565.6 | 0.0001 |
| 400 | 0.68 | 0.983 | 462.0 | 0.0001 |
| 500 | 0.55 | 0.949 | 149.0 | 0.0001 |

1 - df = 1, 8 for all analyses. 2 - slope in units of degrees per second.

**Table 3.** Median Fits by Segment Length - Vertical Signals.

| Segment Length (ms) | Median Slope <sup>2</sup> | Rsqr | F-value <sup>1</sup> | p-value |
| --- | --- | --- | --- | --- |
| 50 | 3.05 | 0.870 | 53.7 | 0.0001 |
| 100 | 1.76 | 0.921 | 93.1 | 0.0001 |
| 200 | 1.15 | 0.955 | 170.9 | 0.0001 |
| 300 | 0.88 | 0.985 | 528.8 | 0.0001 |
| 400 | 0.71 | 0.987 | 586 | 0.0001 |
| 500 | 0.58 | 0.976 | 321.2 | 0.0001 |

1 - df = 1, 8 for all analyses. 2 - slope in units of degrees per second.

### Comparison of Statistical Results from KW Tests across various segment lengths

Once again, we analyzed six different segment lengths (50, 100, 200, 300, 400 and 500 ms). The horizontal position results are presented in Table 4. All of these *χ*^2^ tests were strongly statistically significant, indicating that for all segment lengths, fixation drift slope was strongly related to epoch number. As is evident from this table, the number of fixations included in each slope estimate was huge. The results for vertical position are presented in Table 5. It is noteworthy that all of the *χ*^2^ values for the vertical position signals were much larger than the *χ*^2^ values for the horizontal position signals. Thus, it appears that the statistical evidence for differences in slope estimate across epochs was greater for the vertical signals than the horizontal signals.

**Table 4.** Results of Kruskal-Wallis Test by Segment Length - Horizontal Signals.

| Segment Length (ms) | Number of Fixations | $\chi^2$ | df numerator | df denominator | p-value |
| --- | --- | --- | --- | --- | --- |
| 50 | 323,260 | 575.8 | 9 | 315,636 | 0.0001 |
| 100 | 285,880 | 662.4 | 9 | 278,408 | 0.0001 |
| 200 | 192,800 | 646.5 | 9 | 186,032 | 0.0001 |
| 300 | 168,140 | 537.9 | 9 | 160,962 | 0.0001 |
| 400 | 154,980 | 437.0 | 9 | 146,275 | 0.0001 |
| 500 | 144,390 | 355.7 | 9 | 134,043 | 0.0001 |

**Table 5.** Results of Kruskal-Wallis Test by Segment Length - Vertical Signals.

| Segment Length (ms) | Number of Fixations | ChiSq | df numerator | df denominator | p-value |
| --- | --- | --- | --- | --- | --- |
| 50 | 323,260 | 915.9 | 9 | 315,636 | 0.0001 |
| 100 | 285,880 | 917.8 | 9 | 278,408 | 0.0001 |
| 200 | 192,800 | 740.6 | 9 | 186,032 | 0.0001 |
| 300 | 168,140 | 537.2 | 9 | 160,962 | 0.0001 |
| 400 | 154,980 | 417.9 | 9 | 146,275 | 0.0001 |
| 500 | 144,390 | 336.4 | 9 | 134,043 | 0.0001 |

### Analysis of Slope Changes across Epochs and Sessions - Horizontal and Vertical Position Signals

The median slopes for all epochs and both sessions for a segment length of 300 ms and for horizontal position signals are presented in Fig. 2. Note the linear trend across epochs and sessions. Given the appearance of these results, for analysis of the linear trend, the epochs from session 2 were considered as epochs 6 to 10. The linear relationship of drift to epoch is very strong. This pattern of results suggests that the “time on task” (TOT) effects on drift from session 2 pick up where the TOT drift effects from Session 1 end. This is surprising because there is ≈ 20 minutes on average between session 1 and session 2. Also the random saccade task on which these analyses are based is only 1 of seven tasks in each session. The results for vertical drift are presented in Fig. 3. Examination of the plots for all of the segment lengths analyzed (https://hdl.handle.net/10877/19260) revealed that the results were somewhat less clear for segment lengths of only 50 or 100 ms, but that all of the analyses of segment lengths from 200 to 500 ms were quite clear and similar.

**Fig 2.**
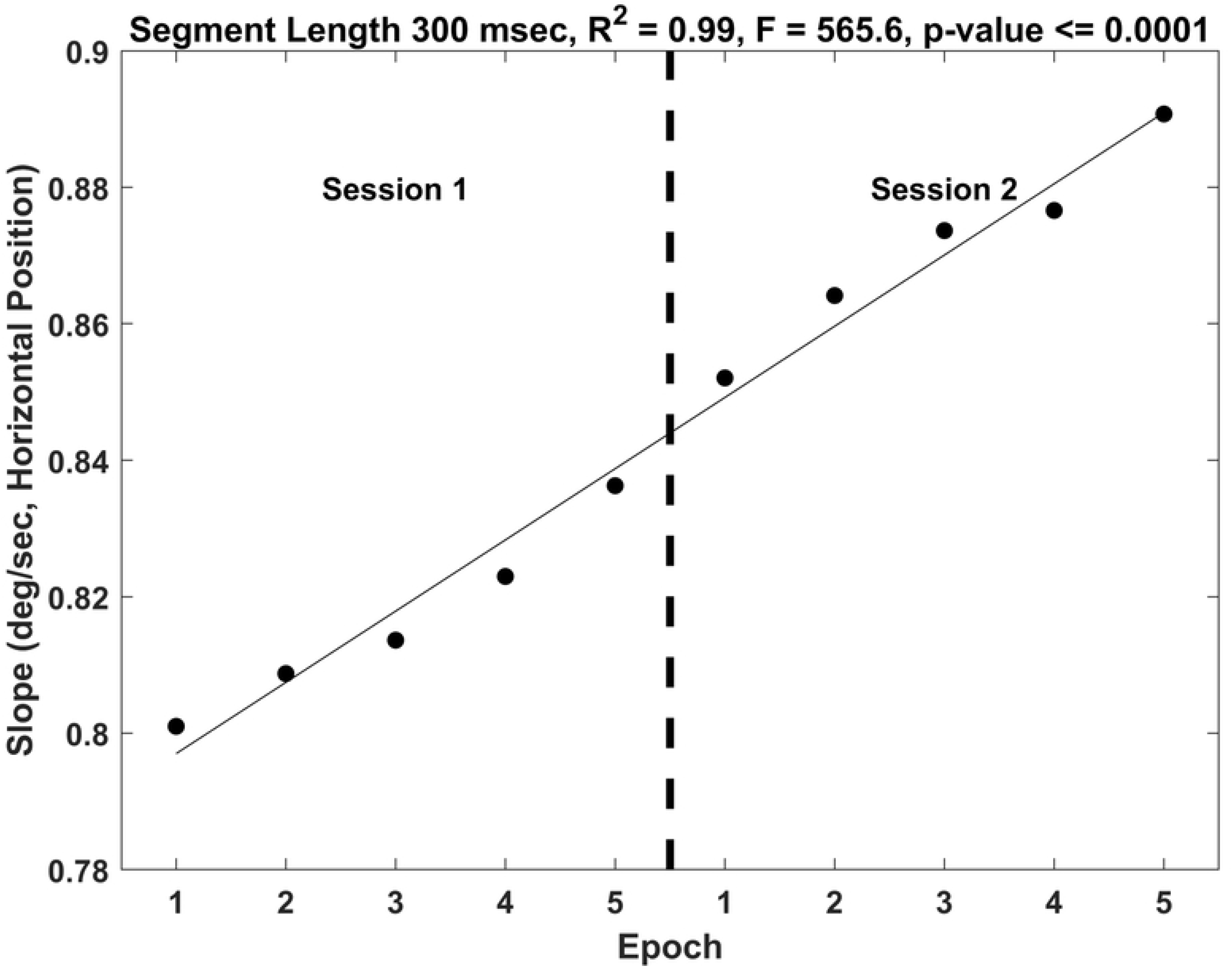
Each dot represents a median slope per epoch. This is for horizontal position signals. Each epoch is 20,000 ms. Also plotted is the regression line relating median slope to epoch, renumbering epochs across sessions 1 and 2 from 1 to 10. Note the almost perfect *R*^2^ of this relationship, the huge F-value and the very significant p-value.

**Fig 3.**
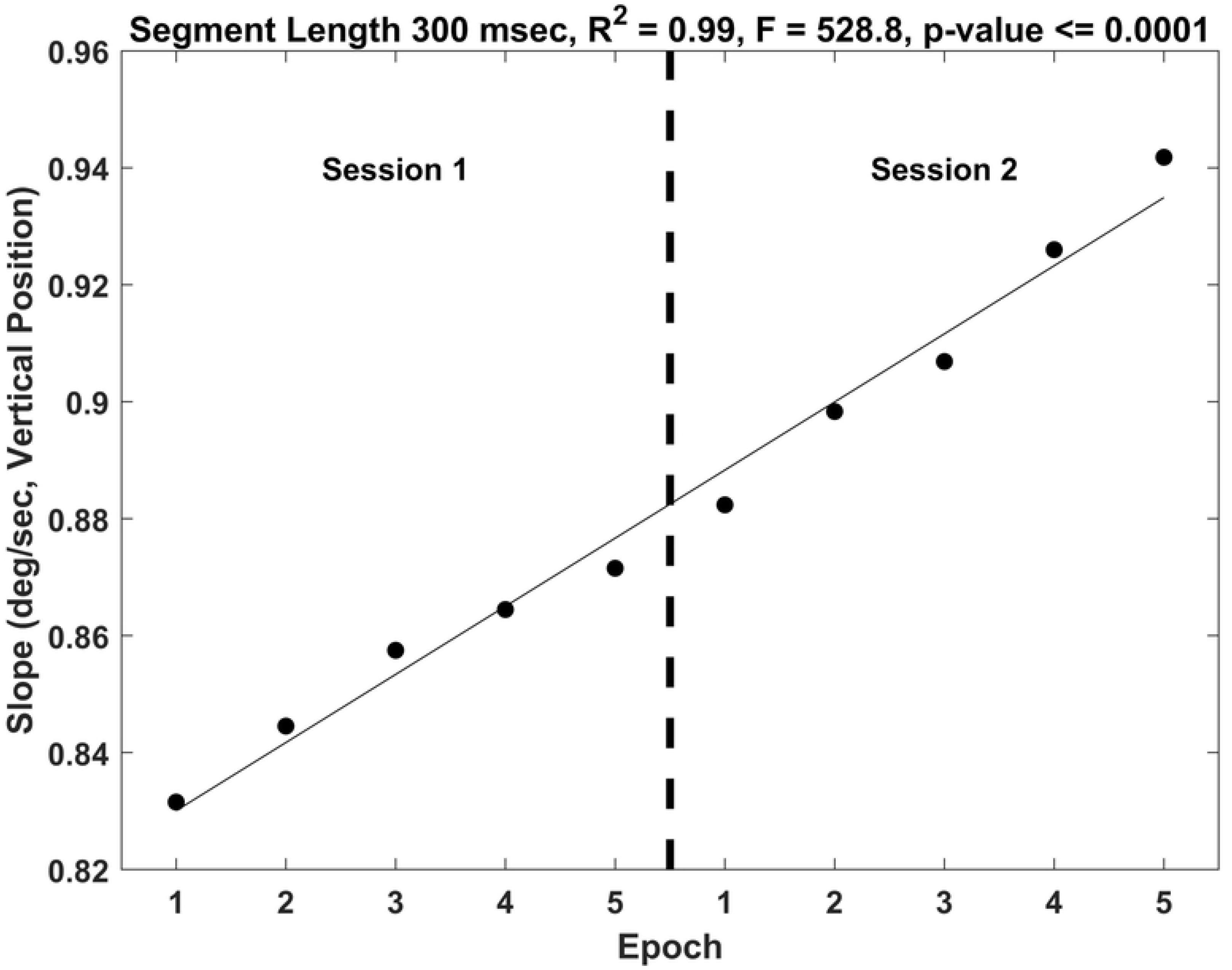
This is the same analysis presented in Fig. 2 for vertical position signals. See caption for Fig. 2.

### Analysis of KW Test Results - Horizontal and Vertical Position Signals

The KW-test answers the following question: is the fixation drift slope significantly different across epochs. The results for horizontal position are presented in Fig. 4. As we already noted above, the *χ*^2^ for this analysis was highly statistically significant. The red dots are mean ranks (the basis of the KW analysis) and the vertical lines indicate statistical significance after controlling for multiple comparisons. Any pair of mean ranks are significantly different if their vertical red lines do not overlap. The results for the vertical position signals are presented in Fig. 5.

**Fig 4.**
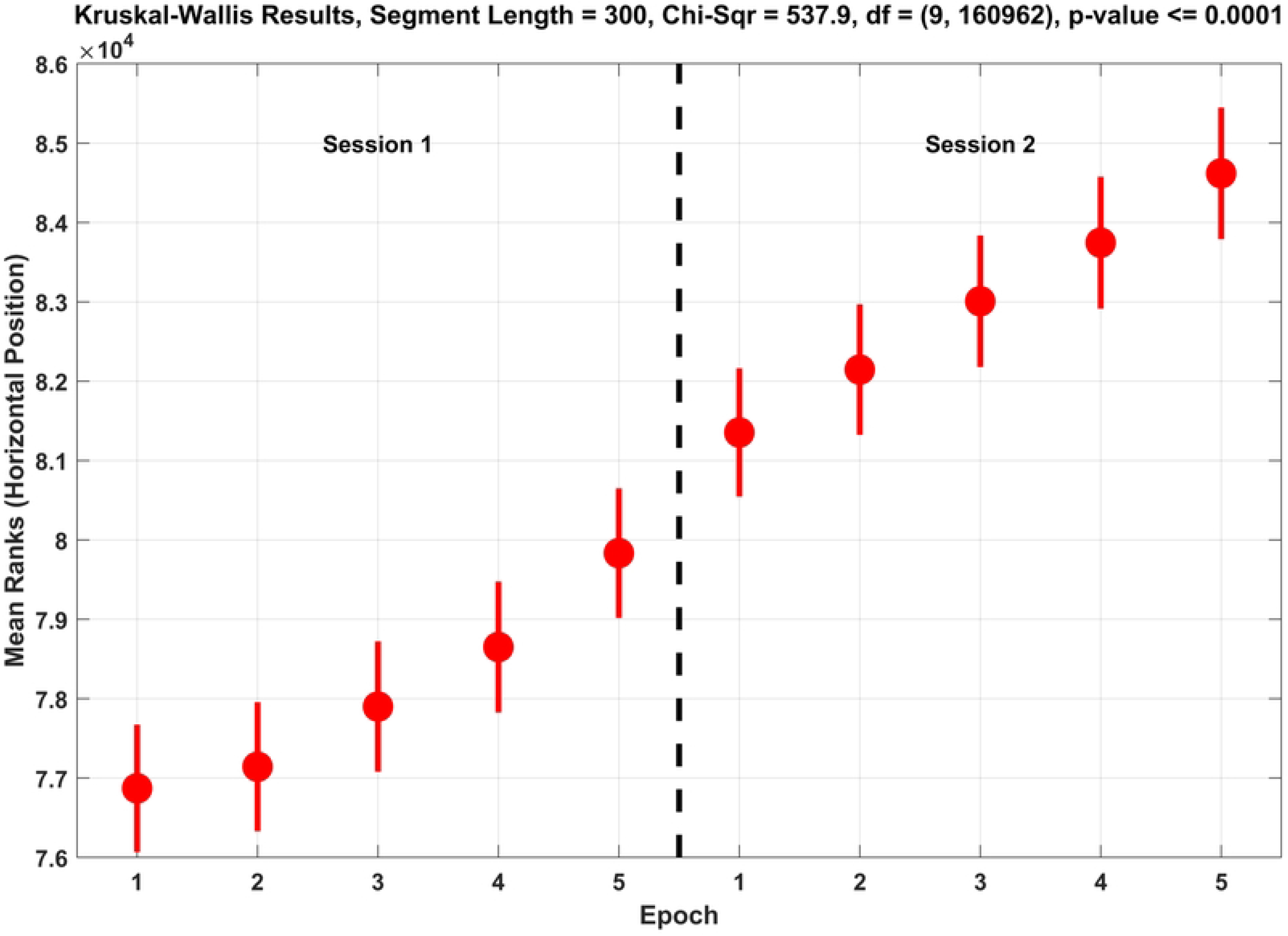
This plot presents the statistical analysis of slopes for horizontal position signals based on a Kruskall-Wallis (K-W) *χ*^2^ test comparing changes across epochs, treating epochs across Sessions 1 and 2 as running from 1 to *10. χ*^2^ = 537.9, *p <* 0.0001. The significant K-W test was followed up with post-hoc multiple comparisons across the epochs. The red circles are the mean ranks for each epoch. Any pair of mean ranks where the red lines do not overlap are statistically significant different (*p <*= 0.05, corrected for multiple comparisons).

**Fig 5.**
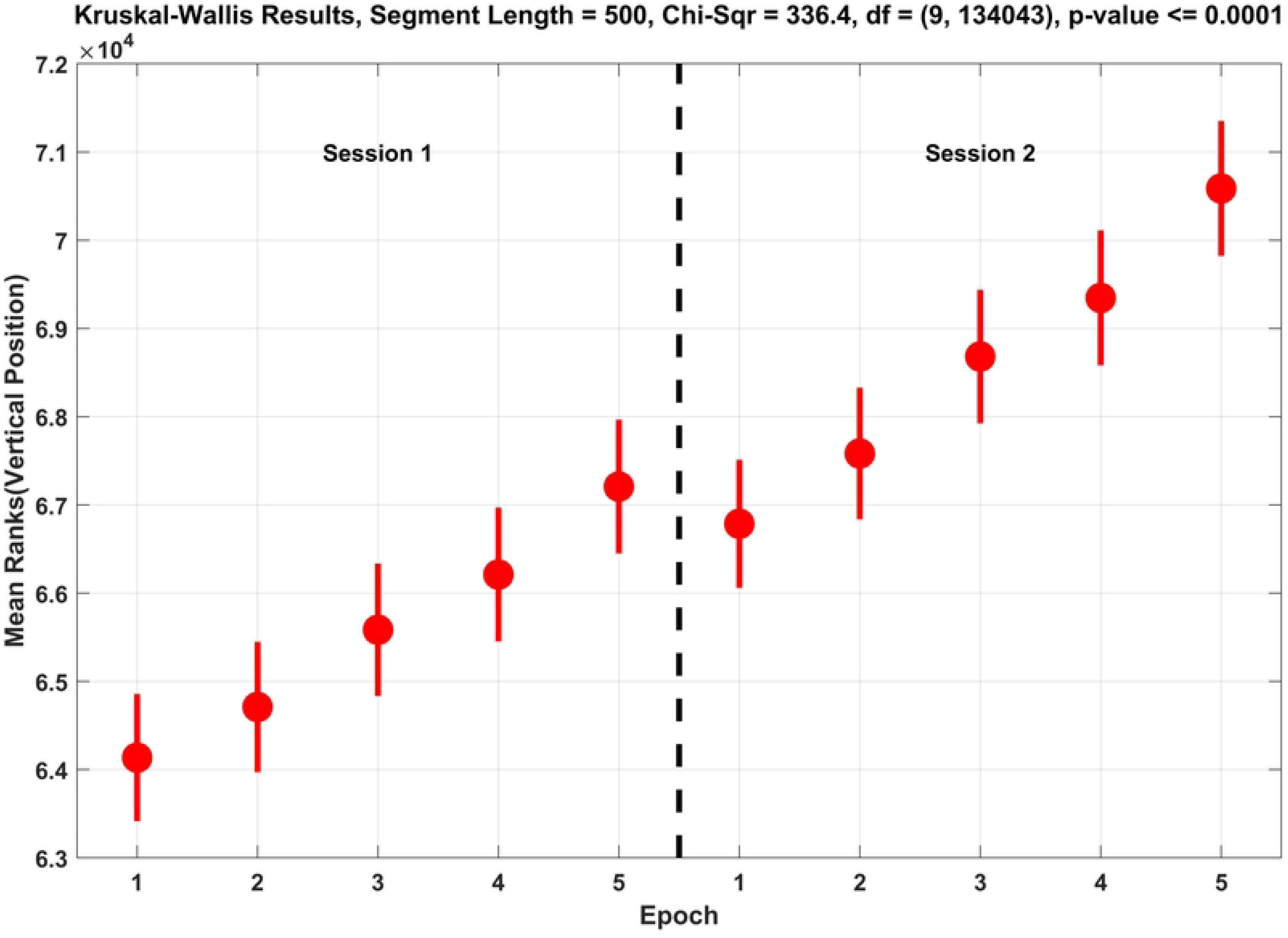
This is the same analysis presented in Fig. 3 for vertical position signals. See caption for Fig. 3.

## Discussion

The main finding of the present study is that the slope of linear fixation drift increased as a function of TOT. Each subject in our study performed the random saccade task twice (two sessions) for each laboratory visit. Although the two sessions were separated by approximately 20 minutes and multiple other eye movement tasks, it was surprising to find that the slope of fixation drift was progressive from one session to the next. Each session was divided into five 20,000 ms epochs. Not only did we find that, within each session, there was a linear trend of increased fixation drift slope across epochs, but, if we renumbered the epochs for session 2 to go from 6 to 10, a linear trend from epoch 1 to epoch 10 was noted. In other words, our results across two 100 second sessions were similar to what we might have expected if we had just one, 200 second task.

Our measure of drift was distinctive from that of the key prior study relating drift velocity to fatigue [4]. In that study, drift was measured as the mean or peak velocity during each fixation. We specifically evaluated *linear* drift. For each fixation, starting with the first sample, we fit a regression to the position data (horizontal or vertical) over several segment lengths. After the results from the first sample were determined, we proceeded to evaluate the slope starting from the second sample, then starting with the third sample, etc… Each fixation was represented by the peak slope across all possible regressions given a particular segment length. These slopes were evaluating linear change over time in degrees per second. Since we were not interested in the direction of the linear drift but only its magnitude, we took the absolute value of the slope from each regression. In our view, the inherent logic of our measures as well as the the pattern and statistical significance of our results justify the use of this measure of linear drift.

We evaluated six different segment lengths (50, 100, 200, 300, 400 and 500 ms). To keep the number of figures in a reasonable range, we presented only the results for 300 ms, but provide a link to figures representing the analyses of all segment lengths. The results for 300 ms were particularly clear, but the same basic pattern of results was consistent across segment lengths.

The Di Stasi, et al. study [4] related their drift findings to fatigue. In that case, the study period was 2 hours. A recent review of mental fatigue using eye metrics found that the minimum study length was 30 min and the median study length was much longer than that. Since our study only covered 200 seconds, we do not refer to our findings as evidence of “fatigue”. Rather, we refer to our findings as related to TOT. However, it seems likely to us that our findings are related to a construct like oculomotor fatigue on a much briefer time scale. However, other changes occur over time such as learning, increasing familiarity with the task, decreases in motivation or increases in boredom cannot be ruled out.

## Conclusion

In conclusion, we found that the slope of linear drift during fixation increases with TOT. This was true regardless of the length of time over which we evaluated drift (50, 100, 200, 300, 400 and 500 ms), although effects for intervals greater than 100 msec were stronger and more consistent. Although we think it is likely that these changes reflect increased oculomotor “fatigue” of a sort over a very short time frame, all we can say for sure is that the findings are related to time-on-task. Other possibilities need to be considered.

## Acknowledgments

We wish to acknowledge our the efforts of our graduate student, Samantha D. Aziz. Sam carefully reviewed our manuscript and provided detailed suggestions and pointed out potential errors or other flaws.

